# Reduced-nutrient leachates in cash cover crop-soybean systems

**DOI:** 10.1101/254169

**Authors:** Matthew D. Thom, Frank Forcella, Carrie A. Eberle, Heather L. Matthees, Sharon L. Weyers, Russ W. Gesch, Matthew A. Ott, Gary W. Feyereisen, Jeffrey S. Strock, Don Wyse

## Abstract

Over-wintering crops are known to reduce nutrients in soil leachate in spring, but little economic incentive is available to grow these crops in the Upper Midwest. New oilseed-bearing cash cover crops, such as winter camelina and pennycress, may provide the needed incentives. However, the abilities of these crops to sequester labile soil nutrients are unknown. We used lysimeters buried at 30, 60, and 100 cm to examine nitrate and soil reactive phosphorus (SRP) in six soybean cropping system treatments: clean till, no-till, and autumn-seeded radish, winter rye, pennycress, and winter camelina. Radish winter-killed naturally, winter rye was killed with a glyphosate, and pennycress and winter camelina were allowed to mature naturally after relay sowing of soybean. Leachate chemistry was studied for the autumn, spring, and summer periods over two growing season. In autumn, leachates under radish and winter rye tended to have the lowest nitrate levels. In spring, differences among nitrate levels across treatments were greater than at any other time period, with values much lower under pennycress and winter camelina treatments than other treatments. In summer, nitrate levels were more uniform, with the lowest values occurring where soybean grew best. In general, cash cover crops like pennycress and winter camelina provide both environmental and economic resources to growers in that they represent cash-generating grain crops that sequester labile soil nutrients, especially in spring, and protect and promote soil health from autumn through early summer.

## Introduction

Conventional cropping systems in the Upper Midwest typically feature fallow soils from October to June (Ochsner et al. 2010; Sindelar et al. 2015). The exposed soil is subject to erosion from wind and water (Skidmore 1988; O’Neal et al. 2005; Li et al. 2007, 2008a; Li et al. 2008b; Sharratt 2011; Palm et al. 2014) and nutrient leaching into subsurface tile drains or groundwater (Strock et al. 2004; Robertson and Vitousek 2009). Leaching of nitrate into tile lines and groundwater is greatest April through June in Minnesota because few plants are growing actively and precipitation predominates over evapotranspiration at that time (Randall et al. 1997). Moreover, 50% of annual drainage occurs by mid-May (Jin and Sands, 2003). Both erosion and leaching result in a depletion of soil organic matter (Pimentel et al. 1995), reduced soil fertility, and a need for increased inputs of fertilizers. Moreover, erosion and leaching contribute to pollution of aquifers, lakes, streams, and entire watersheds (Carpenter et al. 1998, Chambers et al. 2008). Eutrophication of lake and river systems is common in Minnesota and the Upper Midwest, leading to classification of many lake and river systems as unsafe waters (Minnesota Pollution Control Agency 2017).

To combat the problem of pollution from agriculturally intensive areas, over-wintering cover crops are an option that can reduce soil erosion and the pollution of water systems through physical protection of soils and uptake of residual soil nutrients (Dabney et al. 2001; Gesch et al. 2014; Ott et al. 2015; Ott 2018). Analogously, the presence of actively growing plants, such as alfalfa (*Medicago sativa* L.) or grasses and forbs present in conservation reserve plantings (CRP), during April through June nearly eliminates nitrate leaching (Randall et al. 1997). However, some growers may be reluctant to establish CRP or raise alfalfa for economic and logistical reasons (SARE 2015). Accordingly, other over-winter cropping options are needed to keep Minnesota landscapes green in April-June, sequester residual nitrate, reduce erosional losses, and generate profits for growers. New cover crop options exist to maintain green landscapes during critical periods, and these can provide direct incentive or profit by grazing or by maturing for harvest of an additional crop prior to or overlapping with the start of the normal growing season.

The present study evaluated the use of cover crops in an inter-seeded soybean (*Glycine max* [L] Merr.) system to mitigate leaching of N and P from cropping systems. Cover crop treatments included autumn-planted forage radish *(Raphanus sativus* L), winter rye *(Secale cereal* L), field pennycress *(Thlaspi arvense* L), and winter camelina *(Camelina sativa* L), alongside tilled and stubble (no-till) fallow treatments. All treatments were monitored for nutrient leaching. This experimental arrangement allowed us to test the hypothesis that having green cover on the landscape reduces nutrient losses compared to the traditional winter-fallowed soil condition.

## Methods

### Agronomy

This study was conducted from 2014 through 2016 (two complete cropping cycles) on a Barnes loam soil (fine-loamy, mixed, superactive, frigid Calic Hapludoll) at the USDA-ARS Swan Lake Research Farm, Morris, Minnesota (45.68° N, 95.8° W). An automated weather station located nearby recorded all precipitation and temperatures on an hourly basis. Research plots (3.0 x 9.1 m) were arranged on a 2-5% slope in a randomized complete block design including four replicates and two site years (Fig. 1). Plots were located in spring wheat stubble that had been sprayed with 1.1 kg active ingredient ha^−1^ of N-(phosphonomethyl) glycine (glyphosate) at least one week prior to cover crop planting.

**Figure 1.**
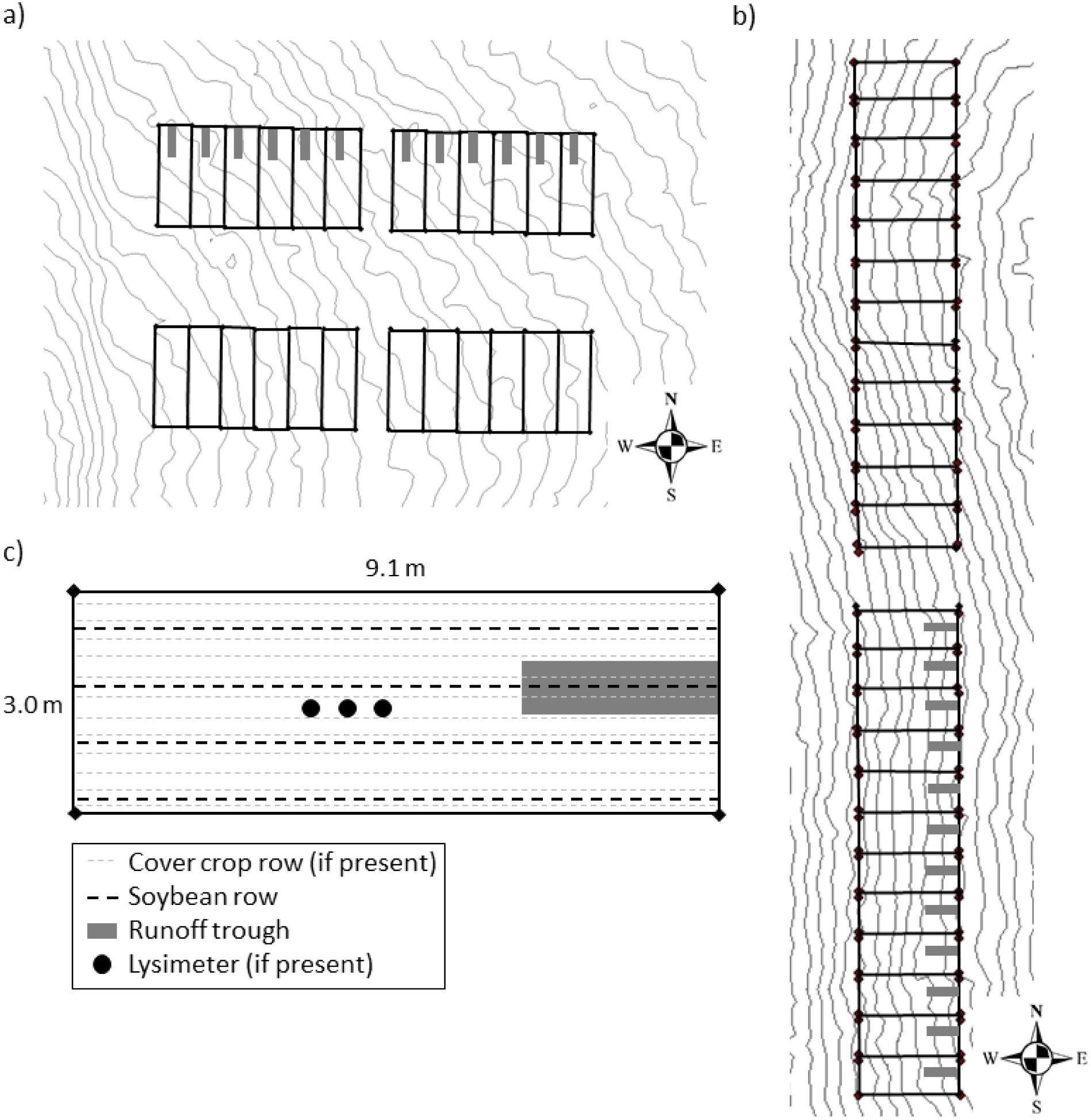
Experimental plots at the Swan Lake Research Farm, Morris, MN. Treatment arrangement and 10 cm topographic contour lines in a) 2014-15 and b) 2015-16; c) Placement of planted crop rows and water sampling equipment

In both 2014 and 2015, plots were prepared in each block with one of six treatments: chisel plow, no-till (wheat stubble), radish, winter rye, pennycress, and winter camelina. Seeding of radish (‘Daikon), winter rye, pennycress (‘Beecher Farms’), and winter camelina (‘Joelle’) cover crops was conducted with a small no-till plot drill (Plotter’s Choice, Kasco Manufacturing, Shelbyville, IN) on 31 August 2014 and 2 September 2015 in 12 rows spaced 25 cm apart at 11, 76, 6.7, and 6.7 kg ha^−1^ and 1.3,1.3, 0.6, and 0.6 cm deep, respectively. The chisel plowing (15-20 cm deep) was conducted on the same day as seeding, and the stubble treatment was left undisturbed.

The following spring, pennycress and winter camelina treatments were fertilized with N-P-K at rates of 80-30-30 kg ha^−1^. Fertilizers were applied as urea, diammonium phosphate, and potash, respectively, and dates were 20 April 2015 and 15 April 2016). The chisel plow treatment was disked and harrowed (30 April 2015 and 22 April 2016), and then all plots were sown with four rows of glyphosate-tolerant soybean (‘Pioneer P09T74R2’) at 445,000 plants ha^−1^ with a drill seeder. Soybean row spacing was 76 cm, and rows were inter-seeded between cover crop rows, if present, to minimize potential crop-to-crop competition. Immediately following soybean planting, winter rye was killed with glyphosate (1 May 2015 and 22 April 2016), but pennycress and winter camelina were allowed to grow. The radish cover crop winter-killed naturally.

Seed harvesting of pennycress occurred on 23 June 23 2015 and 16 June 2016, and that for winter camelina on 2 July 2015 and 24 June 2015). A plot combine (Hege, model 160) was used with the header positioned above the inter-seeded and growing soybean plants in two 1.5 x 10 m areas of each plot. Subsequently, glyphosate was applied as needed for control of weeds in soybean. Soybean seeds were harvested at maturity in late September or early October in a 1.5 x 10 m area in each plot with a plot combine.

### Cover of standing crop

Normalized difference vegetation index (NDVI) was measured weekly using an active canopy sensor (model ACS-470, Holland Scientific, Lincoln, NE) on all plots following emergence of cover crops in the autumn, ceasing after daily maximum temperatures dropped below 10°C. The NDVI measurements were resumed in the spring following the final hard frost (-2°C) and continued until the harvest of soybeans in early autumn. Measurements were taken by holding the sensor approximately 1 m above the top of the plant canopy on the second row of cover crop or soybean. The same row was sampled each time to document the development of green cover of the plot over time. Measurements within the plot and among replicates were averaged to produce an overall measure of green cover for each treatment.

### Water sample collection and analysis

Soil water was sampled using suction cup lysimeters inserted into boreholes placed in the center of the experimental plots using a hydraulic soil probe (Fig. 1c). To ensure optimal soil to lysimeter contact, soil from the bottom of the core was mixed with water to form a slurry, which was placed into the borehole first, followed by insertion of the lysimeters. In 2014-2015, each plot in two blocks was outfitted with two lysimeters, one at 30 cm and the other at 60 cm soil depth. In 2015-2016, each plot in three blocks was outfitted similarly, with a third SCL at 100 cm depth placed in the no-till, winter rye, pennycress, and camelina treatments. Twenty-four hours after all precipitation events of 6 mm or greater, lysimeters were pressurized (vacuumed) to −60 kPa with hand pumps. Water, if present, was collected using hand pumps 24 h after the vacuum was applied to allow time for water to be drawn from the surrounding hemisphere of soil into the ceramic cup at the base of the lysimeter. No samples were collected during winter (December through March).

Collected water samples were taken to the laboratory for processing and chemical analysis. The volume of water collected from the runoff troughs was measured to the nearest ml. Approximately 50 ml of water was filtered with a 25 mm acrylic housed 0.45 μm pore polyethersulfone membrane syringe filter and portioned into two 20-ml subsamples for nitrogen and phosphorus analysis. All nitrogen samples were acidified with 2 μ1 of stock sulfuric acid per 1 ml of filtered sample and held between 2-5 °C until analysis. Phosphorus samples were placed in a −10 °C freezer until analysis.

Nitrogen analysis was conducted at the USDA-ARS North Central Soil Conservation Laboratory in Morris, MN. Nitrate was analyzed using the automated cadmium reduction method, ammonium was analyzed using the automated phenate method (A.P.H.A. 1989), and total nitrogen was measured by electrical conductivity detection using a Lachet IL550 TOC-TN analyzer (Hach, Loveland, CO). Soluble reactive phosphorus (SRP) was analyzed at the Soil and Water Management Research Unit in St. Paul, MN, using the automated ascorbic acid reduction method (A.P.H.A. 1989) with a flow-injection analyzer (Lachat QuikChem 8500, Hach Co.).

### Statistical methods

Nutrient data for the lysimeters were separated into three biologically relevant periods: 1) autumn (from September seeding through November, representing cover crop growth from emergence tofirst hard freeze of −2°C; 2) spring (from snowmelt in April through June, representing spring cover crop growth through oilseed harvest); and 3) summer (from July through September, representing the time of pennycress and winter camelina oilseed harvest through to soybean harvest). These biologically relevant periods were confirmed by linear analysis of the cumulative runoff for each treatment, each site-year growing season, which indicated this ternary distribution.

Nutrient data from lysimeter collections were averaged over all sampling events within each of the three biological periods, and across the two site-years. The non-parametric Kruskal-Wallis test was used to identify significant differences among treatments for concentrations of nitrate, total nitrogen, and SRP gathered from lysimeters. Pairwise comparisons were conducted using the Dunn test with Bonferroni correction if significant differences were found from the Kruskal-Wallis test. Mean separation letters were assigned to treatments to distinguish similar and dissimilar groups using web-based software (Dallal 2000). All statistical tests for nutrients were conducted in R (R Core Team 2015).

## Results

### Weather

Air temperatures during the two years of the study were similar to or slightly higher than 30-year averages for the research site (Fig. 2). In contrast, large differences in seasonal patterns of precipitation occurred between the two years. Autumn and winter precipitation in the 2014-15 season was extremely low compared to both 2015-16 and the 30-year average. During autumn 2014 rain fell in 20 events, with a 13-mrm event maximum. This was substantially less than the total precipitation of 117 mm during the same period in 2015, delivered in 18 events, with a 37-mm event maximum. Spring of 2015 (April through June) was near average with 241 mm precipitation recorded in 32 events (57 mm maximum), in contrast to the spring of 2016 with 148 mm from 31 events (22 mm maximum). The summer (post oilseed harvest period of July through September) was again different between years. The former year, 2015, was near-normal with 212 mm (23 events, 42 mm maximum) of rainfall, whereas 2016 was above normal with 321 mm (28 events, 54 mm maximum) of rainfall in 2016.

**Figure 2.**
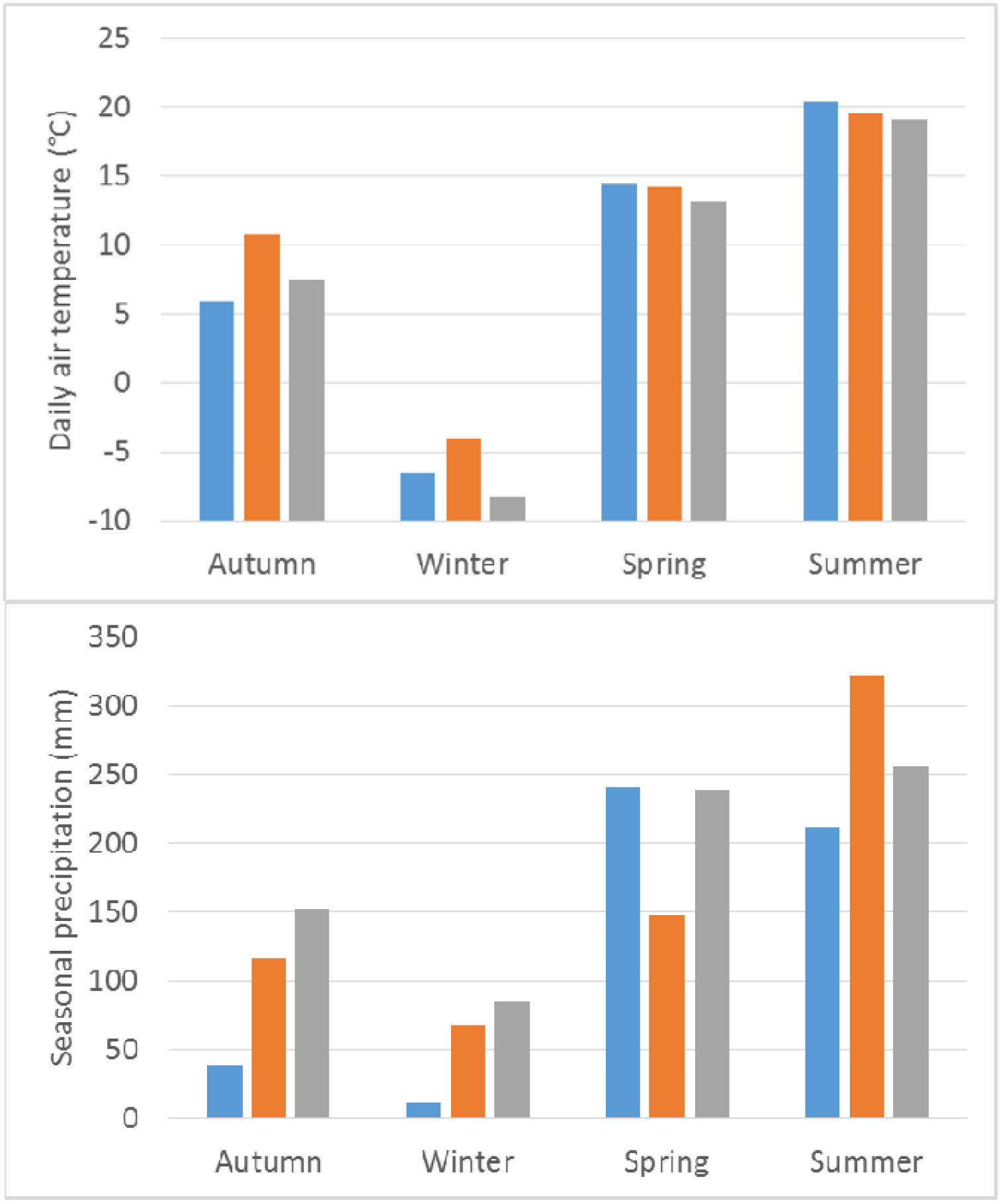
Daily average air temperature (top) and cumulative seasonal precipitation (bottom) during 2014-15 (blue), 2015-16 (orange), and 30-year average (gray) at the Swan Lake Research Farm, Stevens County, Minnesota.

### Green Cover

Measurements of NDVI for each season showed a similar pattern of cover between years, notably in the spring and post-oilseed harvest periods (Fig. 3). In 2014, after autumn establishment, radish and winter rye quickly increased in green cover, with NDVI values exceeding 1. NDVI values of all remaining treatments increased after a brief lag time to > 0.6 (Fig. 3a). This pattern was similar in 2015, but with much lower values, which likely reflected a lack of precipitation until late September to initiate germination of the shallowly sown cover crop seeds (Fig. 3b).

**Figure 3.**
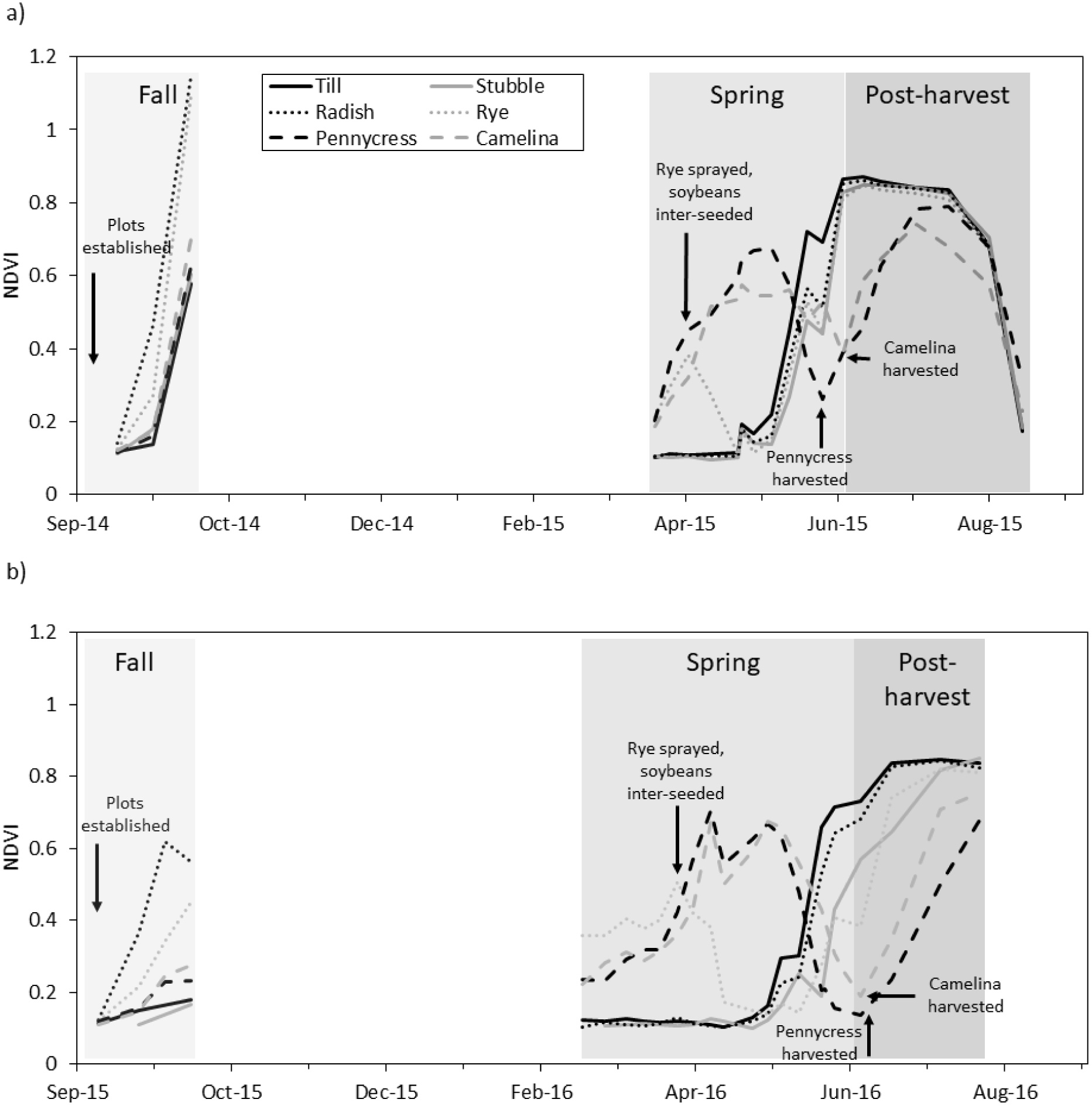
Normalized difference vegetation index (NDVI) for the a) 2014-2015 and b) 2015-2016 growing seasons from experimental plots at the Swan Lake Research Farm, Morris, MN.

For the spring period in both years, stubble, tilled, and radish all remained at the baseline until approximately 30 days after soybeans were inter-seeded into all plots; this was due to winter dieback of seeded or volunteer plants. In both years, winter rye plots increased in green cover until they were terminated with herbicide at soybean inter-seeding, after which time NDVI quickly declined before increasing with cover from soybean emergence and growth roughly 30 days later in 2015 and approximately 45 days later in 2016. NDVI in both pennycress and winter camelina increased at the start of spring measurements, topping at an index value of 0.7 before declining as the crops matured and formed seed heads. Immediately following harvest, the pennycress and winter camelina plots increased in green cover again due to soybean growth, which previously was hindered under the oilseed crop canopies.

### Soil Water Nutrients

For the 2014 autumn period no lysimeter data were collected as precipitation was insufficient to result in soil water saturation. In autumn 2015 the average nitrate and total nitrogen concentrations measured in lysimeter water tended to be lower in radish and winter rye treatments than in the tilled treatment, but not different from concentration in the pennycress and winter camelina treatments (Fig. 4). At 30 cm depth, nitrate in the tilled treatment exceed 50 mg L^−1^. At this same depth, radish, winter rye, pennycress and winter camelina had nitrate values of <5, <5, <20, and <10 mg L^−1^, respectively. No significant differences among treatments occurred for nitrate, or total N at 100 cm, or for SRP at any depth.

**Figure 4.**
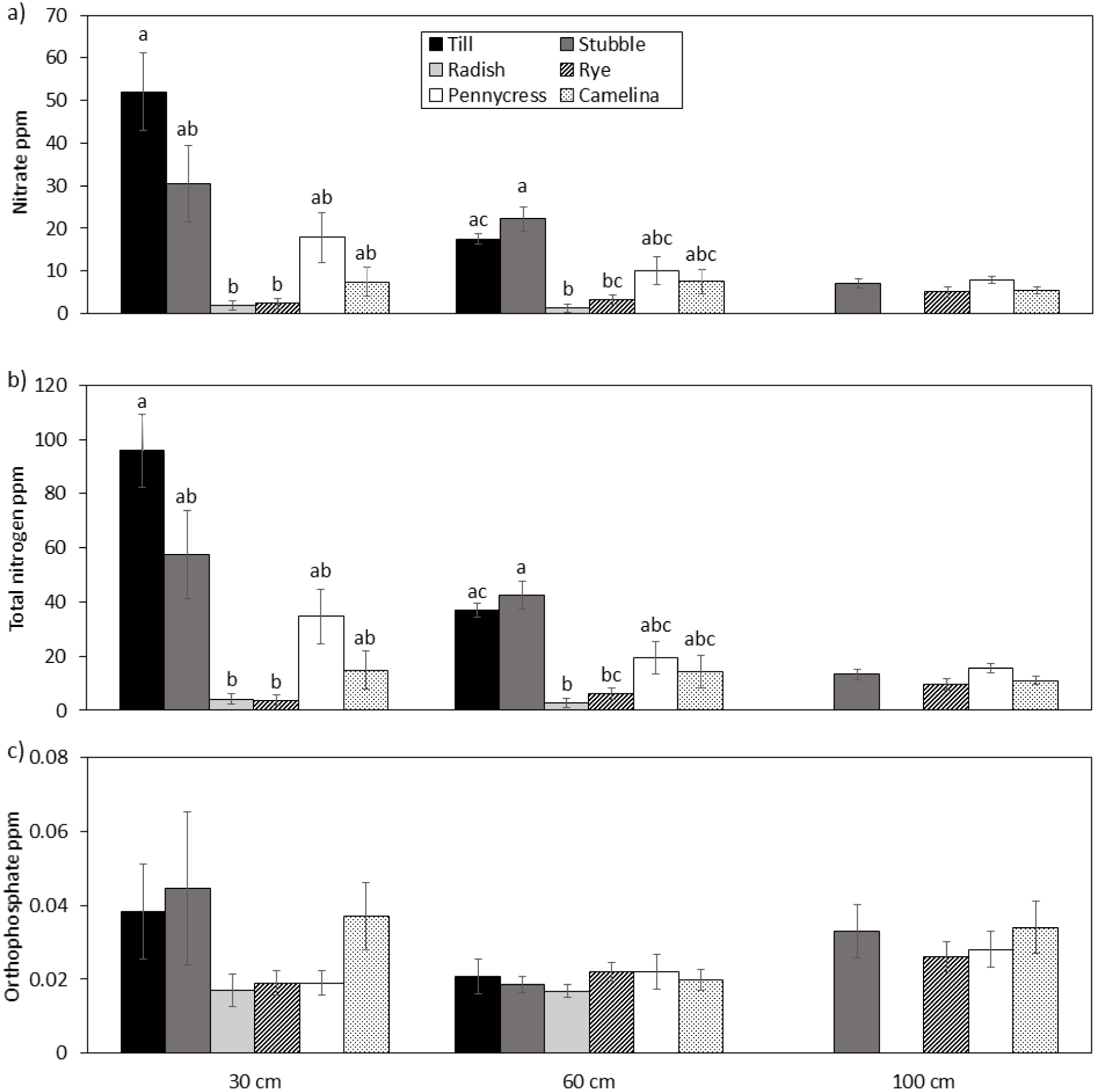
Average autumn (September through November) concentrations (mg L^−1^) of a) nitrate, b) total N and c) SRP (over all sampling dates in 2015-16 growing season) in water samples collected from suction-cup lysimeters installed in experimental plots at the Swan Lake Research Farm, Morris, MN. Columns are means (± SE). Letters represent significant differences (*P*<.05) within a depth, based on the Dunn test. The 100-cm depth lysimeters were installed only in stubble, winter rye, pennycress, and camelina treatments.

For the spring 2015 and 2016 periods, average nitrate and total nitrogen concentrations in soil water from 30 and 60 cm showed very consistent trends with respect to tilled, no-till, and cover crop treatments. For instance, at the 30-cm depth, nitrate concentrations in the tilled treatment > 50 mg L^−1^, whereas concentrations in winter rye, pennycress, and winter camelina all were < 10 mg L^−1^ (Fig. 5). The radish winter-killed and apparently released N to the soil and soil water, which had a nitrate concentration of about 30 mg L^−1^. At 100 cm, no differences were observed for nitrate concentration, but total nitrogen was significantly lower in pennycress, winter camelina and winter rye treatments than in the stubble treatment. No significant differences in SRP among treatments were detected at any depth.

**Figure 5.**
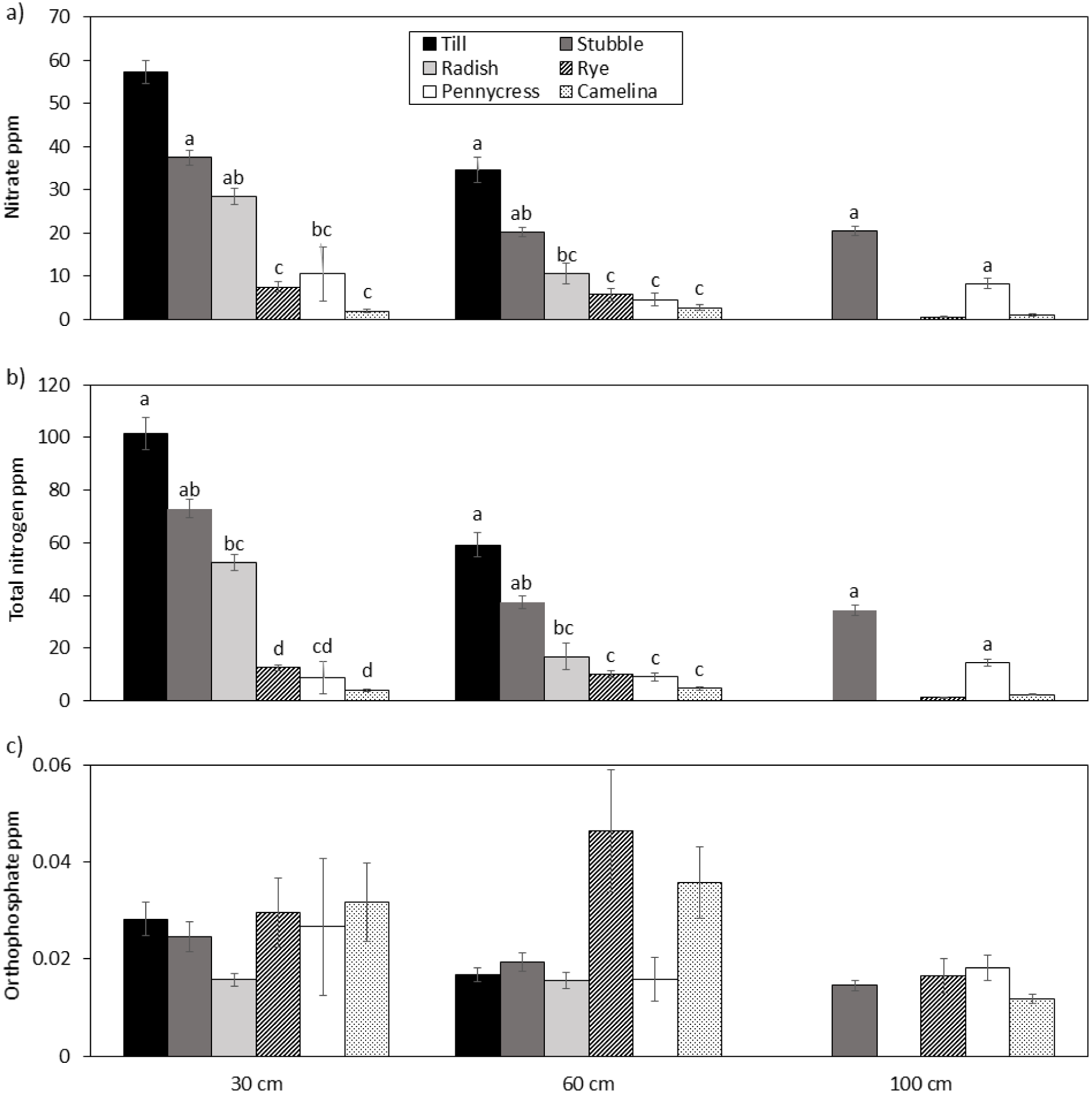
Average spring (April through June) nutrient concentrations (mg L^−1^) of a) nitrate, b) total N, and c) SRP in water samples collected from suction-cup lysimeters installed in experimental plots at the Swan Lake Research Farm, Morris, MN, during the 2014-15 and 2015-16 growing seasons. Columns are pooled means (± SE). Letters represent differences based on the Dunn test. The 100-cm depth lysimeters were installed only in stubble, winter rye, pennycress, and winter camelina in 2015-16.

For the post-harvest periods of 2015 and 2016, average nitrate and total N in leachate from 30 cm and 60 cm depths were much lower than the maximum values seen during autumn or spring (Fig. 6). However, some significant differences still occurred among treatments. The lowest values often were in treatments in which soybean grew best, which tended to be the radish, winter rye, till, and no-till treatments. At 100 cm, average nitrate and total nitrogen values in winter rye were significantly lower than in other treatments. The average SRP was higher in pennycress treatments than some other treatments at both 30 and 60 cm depths (Fig 6c). SRP at 100 cm did not significantly differ among treatments.

**Figure 6.**
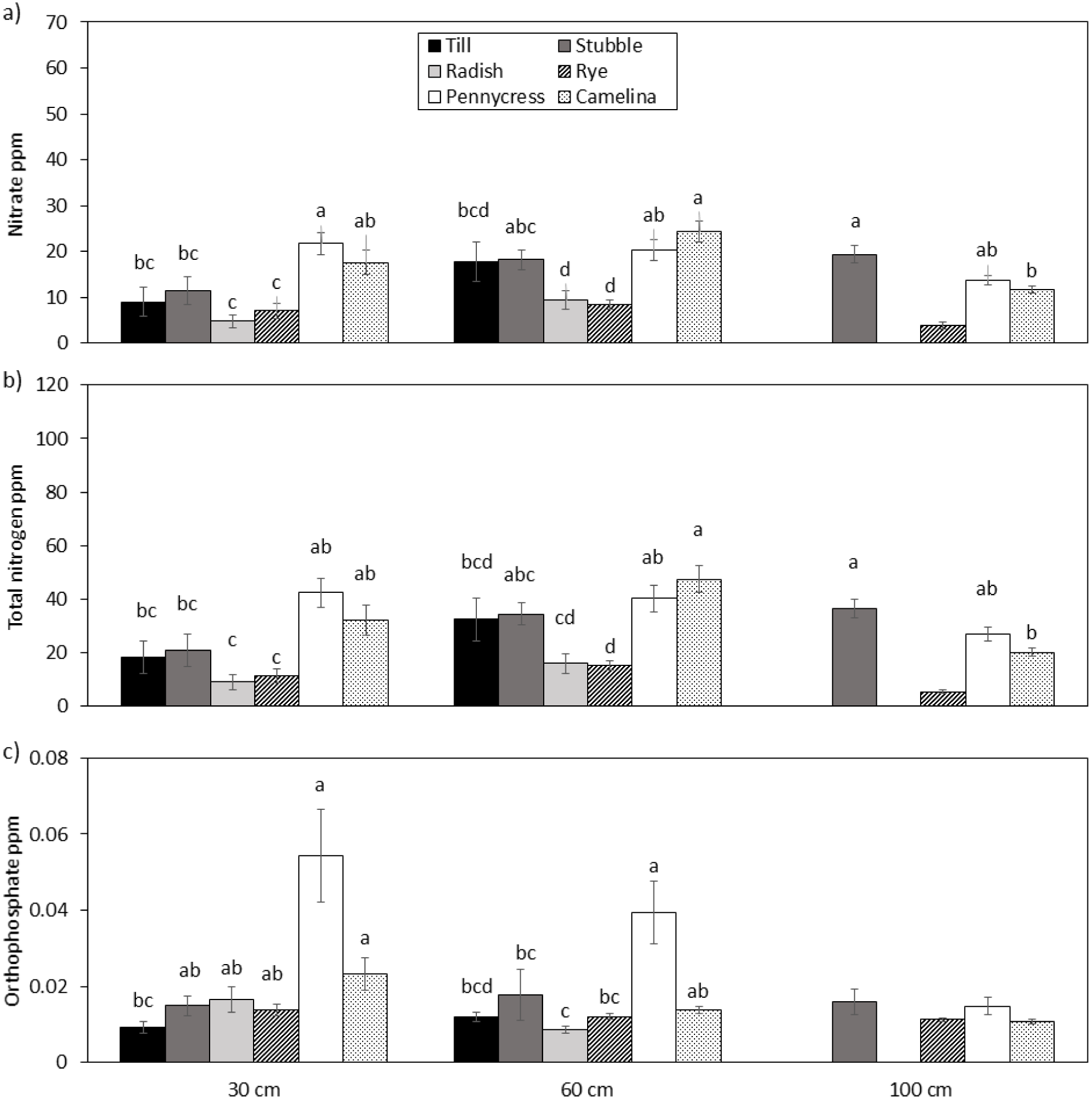
Average summer (post oilseed harvest from July through September) nutrient concentration (mg L^−1^) of a) nitrate, b) total N, and c) SRP in water samples collected from suction-cup lysimeters installed in experimental plots at the Swan Lake Research Farm, Morris, MN, during the 2014-5 and 2015-16 growing seasons. Columns are pooled means (± SE). Letters represent significant differences based on the Dunn test. The 100-cm depth lysimeters were installed only in stubble, winter rye, pennycress, and camelina in 2015-2016.

## Discussion

Results of this study clearly showed that having year-round green cover reduced soil water nutrient concentrations, which might have implications for leaching losses. This was most clear during the spring season with treatments containing the winter annual cover crops winter rye, pennycress, and camelina, compared to plots that were fallow over the winter, including those with radish, which is winter-killed. Traditional winter-fallowed systems had nitrate concentrations in leachate of 20 to 55 mg L^−1^ during spring, whereas analogous concentrations in winter rye, pennycress, and camelina treatments were typically < 10 pp. In the Upper Midwest, spring is a critical time for loss of N and P from agricultural systems due to leaching *via* snowmelt followed by heavy precipitation and low evapotranspiration demand; it is also the season when most nitrate enters groundwater (Randall et al. 1997).

Soybean served the same nutrient-uptake function as the cover crops once this main crop entered the V6-R1 stage, as evidenced by the results of our sampling during the post-oilseed harvest period. In treatments that included inter-seeded soybean with pennycress and winter camelina, there was some degree of continuous green cover over nearly the entire year (i.e., autumn of year 1 to autumn of year 2). Following the harvest of winter oilseeds, soybean cover was lower than in other treatments (Fig. 2; post-harvest period), and it tended to have greater nutrient concentrations in soil water compared to the well-established till, stubble, winter rye, and radish treatments. This greater nutrient concentration was likely related to the lower amount of actively growing soybean cover coupled with relatively high precipitation during the July to August period.

Relay-cropping soybean into standing camelina can reduce soybean biomass and yield (Gesch et al. 2014; Berti et al. 2015), but the total grain produced by both crops usually exceeds that of monoculture soybeans and can provide economic incentive for adoption (Gesch et al. 2014). Nevertheless, the greater N in soil water during the post-harvest period in the pennycress and camelina treatments was much lower (< 20 mg L^−1^) than that found in the leachate of the fallow (i.e., till and stubble) systems during autumn and spring periods.

Cover crops can scavenge residual soil nitrate actively, but uptake depends on climate, soil nitrate levels, planting time, and cover crop species (Gallaher 1977; Delgado et al. 1999; Dabney et al. 2001). In the autumn, radish and winter rye treatments were most efficient at scavenging residual N and reducing soil water N concentrations at both the 30 and 60 cm soil depths compared with pennycress and camelina cover crops. Radish, which does not overwinter in Minnesota, and winter rye, typically have greater biomass in autumn, whereas pennycress and camelina tend to stay in the rosette stage (lower biomass) until spring (Ott 2018). Accordingly, NDVI green cover measurements were greater for radish and winter rye in autumn, indicating greater biomass production, than pennycress and camelina. In the spring, residual N was more effectively scavenged by pennycress, winter camelina and winter rye, despite its April termination, and soil water N concentrations were reduced in all three treatments. In the spring, NDVI data indicated greater green cover and active growth of these oilseed crops. Surprisingly, the radish and stubble treatments, which did not have green cover in the spring prior to soybean planting, also had lower soil water N compared to the tillage treatment. This reduction is likely due to the decomposition of the radish and stubble plant materials resulting in temporary immobilization of N (Kuo et al. 1996; Justes et al. 1999; Kuo and Jellum 2002).

The winter cover crops pennycress and camelina planted in a relay system with soybean reduced soil water nitrogen in comparison to conventionally tilled or other winter fallow scenarios. The improved water quality benefits of pennycress and winter camelina add to their versatility as cover crops in the Upper Midwest, as they can produce a harvestable oilseed for human consumption and industrial use (Zubr 1997; Gesch and Archer 2013) as well as provide floral resources for beneficial insects such as bees (Eberle et al. 2015; Thom et al. 2017). We argue that this combination of economic viability, environmental benefits such as floral resources for pollinators, and potentially reduced nutrient loss provides an important opportunity to increase cropping diversity, improve beneficial insect habitat, and improve impaired water systems that receive drainage from agricultural lands. Incorporating pennycress and winter camelina production into current agricultural practices are not without pitfalls, but the multitude of benefits that these crops are demonstrated to provide are a way of reducing the often harmful yet unintended effects of intensive agricultural production. Careful management of agricultural land to produce increased amounts of food and fuel in an efficient manner is a global priority, one that needs to be solved through innovative and integrated strategies, and the planting of the winter cover crops represents such an approach.

## References

A.P.H.A. 1989. Standard methods for the examination of water and wastewater. American Public Health Association, Washington, DC.

Berti, M., R. Gesch, B. Johnson, Y. Ji, W. Seames, and A. Aponte. 2015. Double‐ and relay-cropping of energy crops in the northern Great Plains, USA. Industrial Crops and Products 75:26–34.

Brady, N. C. and R. R. Weil. 2002. The nature and properties of soils. Prentice Hall, Upper Saddle River, NJ, USA.

Carpenter, S. R., N. F. Caraco, D. L. Cornell, R. W. Howarth, A. N. Sharpley, and V. H. Smith. 1998. Nonpoint pollution of surface waters with phosphorus and nitrogen. Ecol. Appl. 8:559–568.

Chambers, P.A., C. Vis, R.B. Brua, M. Guy, J.M. Culp and G.A. Benoy. 2008. Eutrophication of agricultural streams: Defining nutrient concentrations to protect ecological condition. Water Sci. Technol. 58:2203–2210.

Dabney, S. M., J. A. Delgado, and D. W. Reeves. 2001. Using winter cover crops to improve soil and water quality. Communications in Soil Science and Plant Analysis 32:1221–1250.

Dabney, S. M., R. A. Rebich, and J. W. Pote. 1999. The Mississippi Delta MSEA program. Proceedings of the 10th Conference of the International Soil Conservation Organization, West Lafayette, IN, USA.

Dallal, G. E. 2000. Little Handbook of Statistical Practice. Tufts University, Boston, MA.

Delgado, J. A., R. T. Sparks, R. F. Follett, J. L. Sharkoff, and R. R. Riggenback. 1999. Use of winter cover crops to conserve soil and water quality in the San Luis Valley of South Central Colorado in Lal, and R, eds. Soil quality and soil erosion. CRC Press, Boca Raton, FL, USA.

Eberle, C. A., M. D. Thom, K. T. Nemec, F. Forcella, J. G. Lundgren, R. W. Gesch, W. E. Riedell, S. K. Papiernik, A. Wagner, D. H. Peterson, and J. J. Eklund. 2015. Using pennycress, camelina, and canola cash cover crops to provision pollinators. Industrial Crops and Products 75:20–25.

Gallaher, R. N. 1977. Soil-moisture conservation and yield of crops no-till planted in rye. Soil Science Society of America Journal 41:145–147.

Gerlach, T. 1967. Hillslope troughs for measuring sediment movement. Revue Geomorphologie Dynamique 4:173.

Gesch, R. W. and D. W. Archer. 2013. Double-cropping with winter camelina in the northern Corn Belt to produce fuel and food. Industrial Crops and Products 44:718–725.

Gesch, R. W., D. W. Archer, and M. T. Berti. 2014. Dual Cropping Winter Camelina with Soybean in the Northern Corn Belt. Agronomy Journal 106:1735–1745.

Gu, S., G. Gruau, R. Dupas, C. Rumpel, A. Crème, O. Fovet, C. Gascuel-Odoux, L. Jeanneau, G. Humbert, and P. Petitjean. 2017. Release of dissolved phosphorus from riparian wetlands: Evidence for complex interactions among hydroclimate variability, topography and soil properties. Science of the Total Environment 598:421–431.

Jin, C.-X., G.R. Sands. 2003. The long-term field-scale hydrology of subsurface drainage systems in a cold climate. Trans. ASAE 46:1011–1021.

Justes, E., B. Mary, and B. Nicolardot. 1999. Comparing the effectiveness of radish cover crop, oilseed rape volunteers and oilseed rape residues incorporation for reducing nitrate leaching. Nutrient Cycling in Agroecosystems 55:207–220.

Kinnell, P. I. A. 2016. A review of the design and operation of runoff and soil loss plots. CATENA 145:257–265.

Kruskal, V. J. 1964. Multidimensional scaling by optimizing goodness of fit to a nonmetric hypothesis. Psychometrika 29:1–27.

Kuo, S. and E. J. Jellum. 2002. Influence of winter cover crop and residue management on soil nitrogen availability and corn. Agronomy Journal 94:501–508.

Kuo, S., U. M. Sainju, and E. Jellum. 1996. Winter cover cropping influence on nitrogen mineralization, presidedress soil nitrate test, and corn yields. Biology and Fertility of Soils 22:310–317.

Landsberg, J. H. 2002. The effects of harmful algal blooms on aquatic organisms. Reviews in Fisheries Science 10:113–390.

Li, S., D. A. Lobb, M. J. Lindstrom, and A. Farenhorst. 2007. Tillage and water erosion on different landscapes in the northern North American Great Plains evaluated using 137Cs technique and soil erosion models. CATENA 70:493–505.

Li, S., D. A. Lobb, M. J. Lindstrom, and A. Farenhorst. 2008a. Patterns of water and tillage erosion on topographically complex landscapes in the North American Great Plains. Journal of Soil and Water Conservation 63:37–46.

Li, S., D. A. Lobb, M. J. Lindstrom, S. K. Papiernik, and A. Farenhorst. 2008b. Modeling tillage-induced redistribution of soil mass and its constituents within different landscapes. Soil Science Society of America Journal 72:167–179.

Liu, A., B. L. Ma, and A. A. Bomke. 2005. Effects of cover crops on soil aggregate stability, total organic carbon, and polysaccharides. Soil Sci. Soc. Am. J. 69:2041–2048. doi:10.2136/sssaj2005.0032

Loughran, R. J. 1989. The measurement of soil erosion. Progress in Physical Geography 13:216–233.

Mather, P. M. 1976. Computational methods of multivariate analysis in physical geography. Wiley and Sons, line, London, England.

McCune, B. and M. J. Mefford. 2011. PC-ORD. Multivariate Analysis of Ecological Data. MjM Software, Gleneden Beach, Oregon.

Minnesota Pollution Control Agency. 2017. Minnesota’s Impaired Waters List. https://www.pca.state.mn.us/water/minnesotas-impaired-waters-list (accessed June 13 2017).

O’Neal, M. R., M. A. Nearing, R. C. Vining, J. Southworth, and R. A. Pfeifer. 2005. Climate change impacts on soil erosion in Midwest United States with changes in crop management. CATENA 61:165–184.

Ochsner, T. E., K. A. Albrecht, T. W. Schumacher, J. M. Baker, and R. J. Berkevich. 2010. Water balance and nitrate leaching under corn in kura clover living mulch. Agronomy Journal 102:1169–1178.

Olson, K.R., M. Al-Kaisi, R. Lal, and L.W. Morton. 2017. Soil ecosystem services and intensified cropping systems. Journal of Soil and Water Conservation, 72:64A–69A.

Ott, M., C. A. Eberle, D. L. Wyse, F. Forcella, and R. Gesch. 2015. Improving Minnesota’s water quality with cash cover crops. American Society of Agronomy, Crop Science Society of America, Soil Science Society of America Annual Meeting, Minneapolis, MN, USA.

Ott, M.A. 2018. Four Cover Crops Dual-Cropped with Soybean: Agronomics, Income, and Nutrient Uptake across Minnesota. M.Sc. Thesis, University of Minneosta, St paul. January 2018.

Palm, C, H. Blanco-Canqui, F. DeClerck, L. Gatere, and P. Grace. 2014. Conservation agriculture and ecosystem services: An overview. Agric, Ecosyst. Environ. 187:87–105.

Pimentel, D., C. Harvey, P. Resosudarmo, K. Sinclair, D. Kurz, M. McNair, S. Crist, L. Shpritz, L. Fitton, R. Saffouri, and R. Blair. 1995. Environmental and economic costs of soil erosion and conservation benefits. Science 267:1117–1123.

Randall, G. W., D. R. Huggins, M. P. Russelle, D. J. Fuchs, W. W. Nelson, and J. L. Anderson. 1997. Nitrate losses through subsurface tile drainage in conservation reserve program, alfalfa, and row crop systems. Journal of Environmental Quality 26:1240–1247.

Regvar, M., K. Vogel, N. Irgel, T. Wraber, U. Hildebrandt, P. Wilde, and H. Bothe. 2003. Colonization of pennycresses *(Thlaspi* spp.) of the Brassicaceae by arbuscular mycorrhizal fungi. Journal of plant physiology 160:615–626.

Robertson, G. P. and P. M. Vitousek. 2009. Nitrogen in Agriculture: Balancing the Cost of an Essential Resource. Pp. 97–125. Annual Review of Environment and Resources. Annual Reviews, Palo Alto.

SARE. 2015. 2014–2015 Annual Report: Cover Crop Survey.

Schindler, D. W. 1977. Evolution of phosphorus limitation in lakes. Science 195:260–262.

Sharpley, A. N. 1981. The contribution of phosphorus leached from crop canopy to losses in surface runoff. Journal of Environmental Quality 10:160–165.

Sharpley, A. N., S. C. Chapra, R. Wedepohl, J. T. Sims, T. C. Daniel, and K. R. Reddy. 1994. Managing agricultural phosphorus for protection of surface waters - issues and options. Journal of Environmental Quality 23:437–451.

Sharpley, A. N. and S. J. Smith. 1991. Effects of cover crops on surface water quality in W. L. Hargrove, ed. Cover crops for clean water. Soil and Water Conservation Society, Ankeny, IA, USA.

Sharratt, B. 2011. Size distribution of windblown sediment emitted from agricultural fields in the Columbia Plateau. Soil Science Society of America Journal 75:1054–1060.

Sindelar, A. J., M. R. Schmer, R. W. Gesch, F. Forcella, C. A. Eberle, M. D. Thom, and D. W. Archer. 2015. Winter oilseed production for biofuel in the U.S. Corn Belt: Opportunities and limitations. GCB Bioenergy:n/a-n/a.

Skidmore, E. L. 1988. Wind erosion. Pp. 203–233 in R. Lal, ed. Soil erosion research methods. Soil and Water Conservation Society, Ankeny, Iowa.

Strock, J. S., P. M. Porter, and M. P. Russelle. 2004. Cover cropping to reduce nitrate loss through subsurface drainage in the northern U.S. Corn Belt. Journal of Environmental Quality 33:1010–1016.

R Core Team. 2015. R: A language and environment for statistical computing. R Foundation for Statistical Computing, Vienna, Austria.

Thorn, M. D., C. A. Eberle, F. Forcella, R. Gesch, and S. Weyers. Specialty oilseed crops provide an abundant source of pollen for pollinators and beneficial insects. Journal of Applied Entomology doi: 10.1111/jen. 12401

USEPA. 1990. National Water Quality Inventory. 1988 Report to Congress. EPA-440-4-90-003. USEPA. https://nepis.epa.gov/Exe/ZyPDF.cgi/00001JVE.PDF?Dockey=00001JVE.PDF (accessed 23 June 2017).

Wolf, D., W. Georgic, and H. A. Klaiber. 2017. Reeling in the damages: Harmful algal blooms’ impact on Lake Erie’s recreational fishing industry. Journal of Environmental Management 199:148–157.

Zubr, J. 1997. Oil-seed crop: Camelina sativa. Industrial Crops and Products 6:113–119.

